## Supplementary material for "Categorical representation from sound and sight in the ventral occipito-temporal cortex of sighted and blind": Categories and stimuli description

SI Table 1. Categories and stimuli.

| CATEGORIES | STIMULI |
|---|---|
| BIRDS | Canary<br>Owl<br>Seagull |
| MAMMALS | Dog<br>Donkey<br>Horse |
| HUMAN VOCALIZATIONS* | Woman<br>Man<br>Man |
| HUMAN NON VOCALIZATIONS | Women laughing<br>Man crying<br>Women yawning |
| TOOLS | Hairdryer<br>Saw<br>Toothbrush |
| GRASPABLE OBJECTS | Guitar<br>Keyboard<br>Telephone |
| BIG MECHANICAL OBJECTS | Church-bell<br>Traffic<br>Train |
| ENVIRONMENTAL SCENES | Storm<br>River<br>Wind |

**Neutral faces in the visual experiment*
