## Supplementary material for "Categorical representation from sound and sight in the ventral occipito-temporal cortex of sighted and blind": Characteristics of early blind participants

SI Table 2. Characteristics of early blind participants.

| Subjects | Age(y) | Sex | Residual visual perception | Onset | Cause of blindness |
|---|---|---|---|---|---|
| EB1 | 30 | M | Diffuse light | 0 | Damaged optic nerve |
| EB2 | 33 | F | No | 0 | Congenital cataracts |
| EB3 | 29 | F | Diffuse light | 4 | Retinopathy |
| EB4 | 67 | M | No | 0 | Congenital glaucoma |
| EB5 | 39 | M | Diffuse light | 0 | Retinopathy |
| EB6 | 26 | M | No | 3 | Infection of the eyes |
| EB7 | 34 | F | No | 0 | Microphtalmia |
| EB8 | 28 | M | No | 3 | Retinopathy |
| EB9 | 29 | F | Diffuse light | 0 | Retinopathy |
| EB10 | 43 | M | No | 0 | Retinopathy |
| EB11 | 35 | F | Diffuse light | 0 | Hypoxia |
| EB12 | 36 | M | No | 0 | Hypoxia |
| EB13 | 27 | F | Diffuse light | 4 | Retinopathy |
| EB14 | 29 | F | Diffuse light | 0 | Retinopathy |
| EB15 | 20 | F | No | 0 | Retinopathy |
| EB16 | 34 | F | No | 0 | Hypoxia |
| EB17 | 27 | F | No | 0 | Damaged optic nerve |

*M, male; F, female*
